# Systematic phenotyping and characterization of the 3xTg-AD mouse model of Alzheimer’s Disease

**DOI:** 10.1101/2021.10.01.462640

**Authors:** Dominic I. Javonillo, Kristine M. Tran, Jimmy Phan, Edna Hingco, Enikö A. Kramár, Celia da Cunha, Stefania Forner, Shimako Kawauchi, Jonathan Neumann, Crystal E. Banh, Michelle Huynh, Dina P. Matheos, Narges Rezaie, Joshua A. Alcantara, Ali Mortazavi, Marcelo A. Wood, Andrea J. Tenner, Grant R. MacGregor, Kim N. Green, Frank M. LaFerla

## Abstract

Animal models of disease are valuable resources for investigating pathogenic mechanisms and potential therapeutic interventions. However, for complex disorders such as Alzheimer’s disease (AD), the generation and availability of innumerous distinct animal models present unique challenges to AD researchers and hinder the success of useful therapies. Here, we conducted an in-depth analysis of the 3xTg-AD mouse model of AD across its lifespan to better inform the field of the various pathologies that appear at specific ages, and comment on drift that has occurred in the development of pathology in this line since its development 20 years ago. This modern characterization of the 3xTg-AD model includes an assessment of impairments in behavior, cognition, and long-term potentiation followed by quantification of amyloid beta (Aβ) plaque burden and neurofibrillary tau tangles, biochemical levels of Aβ and tau protein, and neuropathological markers such as gliosis and accumulation of dystrophic neurites. We also present a novel comparison of the 3xTg-AD model with the 5xFAD model using the same deep-phenotyping characterization pipeline. The results from these analyses are freely available via the AD Knowledge Portal (https://admodelexplorer.synapse.org). Our work demonstrates the utility of a characterization pipeline that generates robust and standardized information relevant to investigating and comparing disease etiologies of current and future models of AD.

**Contribution to the Field Statement:** Alzheimer’s Disease (AD) is an age-related neurodegenerative disorder characterized by progressive memory impairments and affects more than 30 million individuals worldwide. Using animal models of AD, researchers have elucidated disease progression and hallmark pathologies that may underpin the memory impairments seen in patients. However, therapeutic targets have failed to translate successfully from animal studies to human clinical trials, prompting a reassessment of the development, use, and interpretation of data acquired using the innumerous AD animal models available to researchers. To address these shortcomings, we have developed a robust and reproducible modern characterization of pathologies within current and future animal models of AD to better assess distinct pathologies that arise at specific brain regions and ages of different models. Using the popular 3xTg-AD mouse, we demonstrate the utility of these deep-phenotyping analyses and highlight the drift that affected development of pathologies in this line over the past two decades. Utilizing this same systematic characterization, we also perform a direct comparison with 5xFAD mice, another popular animal model of AD. The robust and standardized data generated from these systematic deep-phenotyping analyses are available for broad use by the AD research community to assess, compare, and determine appropriate animal models of AD.

## Introduction

Alzheimer’s Disease (AD) is an age-related neurodegenerative disorder characterized by progressive memory deficits that affects more than 6 million Americans and more than 30 million individuals worldwide (2021). Currently, no cure exists and any therapies are palliative in nature with limited benefit to disease progression or pathogenesis. The two hallmark pathologies observed in the AD brain are extracellular plaques composed mainly of the amyloid-beta peptide (Aβ), and intraneuronal neurofibrillary tangles (NFT’s), composed primarily of hyperphosphorylated tau protein (DeTure and Dickson, 2019). It is not yet fully understood why these pathologies arise in the aged brain and identifying the underlying and likely multiple mechanism(s) for their development remains a key goal. The accumulation of plaques and NFT’s is associated with a cascade of events that ultimately results in extensive synaptic and neuronal loss, which is thought to underlie the memory deficits observed in patients. Neuroimaging and clinical data point to the appearance of cortical plaques as an early predictor of AD, followed by NFT’s, brain atrophy, and then clinically detectable impairments (Soria Lopez et al., 2019). Notably, pathology initiates in discrete brain regions and then spreads, with plaques first appearing in the cortex, while NFT’s initially accumulate in the entorhinal cortex and hippocampus.

Animal models of AD are valuable resources to investigate mechanisms of AD pathogenesis. Indeed, many observations made using mouse models of AD recapitulate those found in human tissue or imaging experiments, and *vice versa*. However, successful therapies developed and tested in these animal models have been universally unsuccessful in human clinical trials, prompting a reassessment of the development, use and interpretations of data acquired from such models (Dallemagne and Rochais, 2020); (Veening-Griffioen et al., 2019). Several issues warrant consideration. First, the vast majority of animal models are based on familial forms of AD, which are rare and inherited in an autosomal dominant fashion, rather than sporadic/late-onset AD which comprises ∼98% of AD cases (Drummond and Wisniewski, 2017). Second, most animal models develop only one of the two hallmark pathologies – either plaques or tangles - while in humans both are required for a transition to AD (Myers and McGonigle, 2019). Third, and perhaps most striking, existing rodent models that produce abundant AD-related pathologies fail to develop the stark synaptic and neuronal loss observed in clinical AD patients (Drummond and Wisniewski, 2017).

To begin to address these shortcomings as well as meeting the critical need for new animal models of late-onset AD (LOAD), the NIH/NIA established Model Organism Development and Evaluation of Late-onset Alzheimer’s Disease (MODEL-AD https://www.model-ad.org/). A key goal of MODEL-AD is to develop new strains of genetically modified mice that have combinations of genetic variants associated with increased risk for LOAD in humans. Doing so will allow investigation of the relative contribution of LOAD genetic risk variants to development of AD-like pathology in mice, as well as informing as to the pathways and mechanisms that lead to development of LOAD. While better recapitulating LOAD pathogenesis, such mouse models should ideally improve translatability across preclinical testing pipelines (Oblak et al., 2020). An early objective of MODEL-AD is to establish phenotyping pipelines for robust and reproducible analysis of the newly developed mouse models of LOAD at independent sites. For an initial test of one such pipeline, we are performing an in-depth phenotyping of two popular animal models of AD; 5xFAD (Oakley et al., 2006), (Forner et al., 2021) and 3xTg-AD mice (Oddo et al., 2003). In addition to validation of the phenotyping pipeline, the results of these analyses will better inform the AD field about the various pathologies that arise, and at which ages and in which brain regions, such that investigators can select the most appropriate model for their hypothesis/experiment. Here, we report the results of our in-depth analysis of the 3xTg-AD model.

The 3xTg-AD mouse (Tg(APPSwe, tauP301L)1Lfa *Psen1*^*tm1Mpm*^/Mmjax) was developed in 2003 and features three familial AD mutations: the Swedish *APP* mutation (KM670/671NL), the *PSEN1* M146V mutation, and the *MAPT* P301L mutation (Oddo et al., 2003). Expression of the multi-copy human *APP* and *MAPT* (TAU) transgenes are regulated by a mouse *Thy1* minigene (Caroni, 1997) while expression of mouse *Psen1* with the M146V mutation is controlled by the cognate mouse *Psen1* locus (Guo et al., 1999). 3xTg-AD mice are usually utilized as homozygous for both the transgene insert and the *Psen1* M146V familial AD mutation. The initial report of the 3xTg-AD model described development of age-related and progressive amyloid and tau pathologies, with extracellular plaques first appearing at 6-months of age followed by neurofibrillary tangles becoming apparent at 12-months of age (Oddo et al., 2003). Additionally, 3xTg-AD mice have also displayed localized neurodegeneration, synaptic impairment, and cognitive deficits at 6 months of age (Drummond and Wisniewski, 2017). The attractive combination of both plaque and tangle development makes the 3xTg-AD mouse regarded as a complete transgenic mouse model of AD pathology (Myers and McGonigle, 2019). Since its original characterization nearly two decades ago, drift has occurred in the phenotypes seen in these mice. This might have occurred due to segregation of alleles in the mixed strain background (combination of C57BL/6, 129/X1 and 129S1), or changes in the transgene copy number within the single site of integration (Goodwin et al., 2019), or other factors. As one of the most widely used animal models in AD studies, we have used the 3xTg-AD mouse as a reference model for the deep-phenotyping pipeline of MODEL-AD to characterize changes in behavior and cognition, long-term potentiation (LTP), neuropathology, and biochemistry across its lifespan (4, 12, and 18 months of age), and to explore the pathology within the current model compared to our original report. We have also conducted a direct comparison with the 5xFAD model using the same deep-phenotyping pipeline. The results of these systematic phenotyping analyses are freely available via the AD Knowledge Portal (https://admodelexplorer.synapse.org) and should be of broad use to the AD scientific AD research community. They also demonstrate the utility of the phenotyping pipeline in providing robust and standardized information relevant to assessing LOAD etiology within current and future models of AD.

## Material and Methods

### Animals

All experiments involving mice were approved by the UC Irvine Institutional Animal Care and Use Committee and were conducted in compliance with all relevant ethical regulations for animal testing and research. All experiments involving mice comply with the Animal Research: Reporting of *In Vivo* Experiments (ARRIVE) guidelines, which are specifically addressed in the supplementary materials.

### Environmental conditions

Animals were housed in autoclaved individual ventilated cages (SuperMouse 750, Lab Products, Seaford, DE) containing autoclaved corncob bedding (Envigo 7092BK 1/8” Teklad, Placentia, CA) and two autoclaved 2” square cotton nestlets (Ancare, Bellmore, NY) plus a LifeSpan multi-level environmental enrichment platform. Tap water (acidified to pH2.5-3.0 with HCl then autoclaved) and food (LabDiet Mouse Irr 6F; LabDiet, St. Louis, MO) were provided *ad libitum*. Cages were changed every 2 weeks with a maximum of 5 adult animals per cage. Room temperature was maintained at 72 ± 2°F, with ambient room humidity (average 40-60% RH, range 10 - 70%). Light cycle was 14h light / 10h dark, lights on at 06.30h and off at 20.30h.

### Mice

Homozygous 3xTg-AD (B6;129-Tg(APPSwe, tauP301L)1Lfa *Psen1*^*tm1Mpm*^/Mmjax mice were obtained from a closed colony maintained in the laboratory of F.M.L.. In 2018, this stock was genotyped using SNP markers and shown to be homozygous for 129×1/129S1 alleles at ∼35% of the genome, homozygous for C57BL/6 alleles at ∼ 50% of the genome and heterozygous for alleles of 129×1/129S1 and C57BL/6 at ∼ 15% of the genome. At that time, qPCR analysis of the APPSwe and TAUP301L cDNA’s in 3xTg-AD mice from the LaFerla colony and those from Jackson Laboratory (Stock # 34830) showed a similar relative copy number for each cDNA. Experimental and control mice for this study were generated as follows. First, sperm from 3xTg-AD homozygous animals from the LaFerla colony was used to fertilize oocytes from B6129SF2/J mice (Jackson Laboratory, Stock # 101045) and zygotes were transferred to pseudopregnant dams. F1 offspring were genotyped to verify heterozygosity for the co-integrated *Thy1-APPSwe* and *Thy1-TAUP301L* transgene array at ∼ 87.9Mb on chromosome 2, and the I145V/M146V mutations in *Psen1* on chromosome 12. F1 heterozygous mice were intercrossed to generate F2 offspring that were genotyped to identify animals homozygous for both the transgene array and *Psen1* mutations, (i.e. experimental 3xTg-AD homozygotes) or *Psen1*^*+/+*^ and non-transgenic (i.e. wild-type control). For simplicity, throughout the text, 3xTg-AD homozygous animals are referred to as “3xTg-AD” and closely related non-transgenic and *Psen* +/+ mice are referred to as “wildtype” (WT) controls. Natural breeding or IVF using gametes from F2 3xTg-AD or WT control mice were used to produce F3 and later generations of 3xTg-AD and WT control mice for experimental analysis. All animals were generated by the Transgenic Mouse Facility at UCI.

### Genotyping

To genotype for the presence of the transgene array and to monitor the relative transgene copy number over each generation, we used hydrolysis probes that hybridize to the APP(Swe) mutation (For 5’-TGGGTTCAAACAAAGGTGCAA -3’, Rev 5’-GATGACGATCACTGTCGCTATGAC-3’, Probe 5’-CATTGGACTCATGGTGGGCGGTG-3’) and hTAU (P301L) mutation (For 5’-GCGGGAAGGTGCAGATAAT-3’, Rev 5’-CTCCCAGGACGTGTTTGATATT-3’, Probe 5’-CCAGTCCAAGTGTGGCTCAAAGGA-3’). To normalize Ct values, the APP and TAU signals were normalized to a signal from amplification of the mouse *ApoB* locus (For 5’-CACGTGGGCTCCAGCATT-3’, Rev 5’-TCACCAGTCATTTCTGCCTTTG-3’, Probe 5’-CCAATGGTCGGGCACTGCTCAA-3’). To genotype the *Psen1* mutant allele, we used hydrolysis probes to discriminate the WT or mutant *Psen1* allele from the same amplicon (For 5’-CACCCCATTCACAGAAGACA-3’, Rev 5’-CAACCCATAGGCAGGTCAAG-3’ *Psen1* WT probe 5’-TGTCATTGTCATTATGACCATCCT-3’, *Psen1* mutant probe 5’-TCATTGTCGTGGTGACCATC-3’)

### Behavioral Testing

Noldus Ethovision software (Wageningen, Netherlands) was used to video-record and track animal behavior, while analyses were obtained via Ethovision software. The following behavioral paradigms were performed according to prior behavior protocols established by MODEL-AD (Forner et al., 2020), and described in brief as follows.

#### Elevated Plus Maze (EPM)

Mice were placed in the center of an elevated plus maze (arms 6.2 × 75 cm, with side walls 20 cm high on two closed arms, elevated 63 cm above the ground) for five min to assess anxiety. Automated scoring assessed the amount of time each mouse spent cumulatively in the open and closed arms of the maze.

#### Y-maze

Mice were placed facing the back wall of one arm of the Y-maze and video recorded for 8 min to assess short-term memory. An arm entry is defined as the animal’s center point within an arm. Using Noldum EthoVision XT 14 software, the number of alternations was determined by the animal’s entry into all three arms during consecutive choices.

#### Open Field (OF)

Mice were placed in a white box (33.7 × 27.3 × 21.6 cm) for 5 min to assess motor function and anxiety while video recorded for 5 min. Videos were scored for the percentage of time mice spent in the center of arena, distance traveled, and velocity.

#### Contextual Fear Conditioning (CFC)

Behavior was scored using Noldus Ethovision v.14.0.1322. Activity Analysis to detect activity levels and freezing behaviors for both training and testing sessions. Each of the four CFC chambers (Ugo Basile, Germany) are inside a sound-attenuating box with ventilating fans, a dual (visible/I.R.) light, a speaker, and a USB-camera. Each FC-unit has an individual controller on-board. The CFC chamber is cleaned at the start of testing and between every mouse with 70% ethanol and paper towels to eliminate olfactory cues. In the training trial, each mouse is placed in the chamber for 2 min to allow for habituation and exploration of the context, after which a shock is applied for 3 sec at 0.5 mA. The mice are returned to their cages after 30 sec. Twenty-four hr later, testing was conducted, whereby animals were placed in the chamber to explore for 5 min. Sessions are recorded and immobility time was determined using Noldus EthoVision XT 14 software.

### Hippocampal slice preparation and LTP recording

Hippocampal slices were prepared from male and female 3xTg-AD (5 females and 5 males) and WT (5 females and 5 males) mice at 4, 12 and 18 months of age. Following isoflurane anesthesia, mice were decapitated, and the brain was quickly removed and submerged in ice-cold, oxygenated dissection medium containing (in mM): 124 NaCl, 3 KCl, 1.25 KH_2_PO_4_, 5 MgSO_4_, 0 CaCl_2_, 26 NaHCO_3_, and 10 glucose. Coronal hippocampal slices (340 µm) were prepared using a Leica vibrating tissue slicer (Model: VT1000S) before being transferred to an interface recording containing preheated artificial cerebrospinal fluid (aCSF) of the following composition (in mM): 124 NaCl, 3 KCl, 1.25 KH_2_PO_4_, 1.5 MgSO _4_, 2.5 CaCl _2_, 26 NaHCO _3_, and 10 glucose and maintained at 31 ± 1°C. Slices were continuously perfused with this solution at a rate of 1.75-2 ml/min while the surface of the slices were exposed to warm, humidified 95% O_2_ / 5% CO_2_. Recordings began following at least 2 hr of incubation.

Field excitatory postsynaptic potentials (fEPSPs) were recorded from CA1b stratum radiatum using a single glass pipette filled with 2M NaCl (2-3 MΩ) in response to orthodromic stimulation (twisted nichrome wire, 65 µm diameter) of Schaffer collateral-commissural projections in CA1 stratum radiatum. Pulses were administered at 0.05 Hz using a current that elicited a 50% maximal response. Paired-pulse facilitation was measured at 40, 100, and 200 sec intervals prior to setting baseline. After establishing a 10-20 min stable baseline, long-term potentiation (LTP) was induced by delivering 5 ‘theta’ bursts, with each burst consisting of four pulses at 100 Hz and the bursts themselves separated by 200 msec (i.e., theta burst stimulation or TBS). The stimulation intensity was not increased during TBS. Data were collected and digitized by NAC 2.0 Neurodata Acquisition System (Theta Burst Corp., Irvine, CA) and stored on a disk.

### Tissue Collection and Histology Preparation

Mice (n=6 per genotype/age/sex) were euthanized at 4, 12, and 18 months via CO_2_ inhalation and transcardially perfused with 1X phosphate buffered saline (PBS; Sigma-Aldrich, St. Louis, MO). For all subsequent analyses, brains were removed with hemispheres separated along the midline. Brain halves were either drop-fixed in phosphate buffered 4% paraformaldehyde 4°C (Thermo Fisher Scientific, Waltham, MA) for 24 hours at for immunohistochemical staining or micro-dissected (cortex, hippocampus, midbrain) and flash frozen for biochemical analysis. Fixed half brains were coronally sliced at 40 µm using a Leica SM2000R freezing microtome. All brain hemispheres were processed and representative slices (between -2.78mm posterior and –3.38mm posterior to Bregma according to the Allen Mouse Brain Atlas, Reference Atlas version 1, 2008) containing hippocampal and cortical regions from each mouse were used for histological staining and stored in 4°C in cryoprotectant.

### Immunofluorescence Staining

A representative slice from each mouse was selected and pooled together in wells containing slices from mice of the same experimental group (e.g. identical genotype, age, and sex) for each combination of immunofluorescence stains. Unless specified, all stains were performed at 20°C. For Thioflavin-S (Thio-S) staining, free-floating sections were washed with 1X PBS three times (1×10 min, 2×5 min) and incubated for 10 min in 0.5% Thio-S (T1892; Sigma-Aldrich, St. Louis, MO) diluted in 50% ethanol. From this point onward, stained sections were kept in the dark. Sections were rinsed with two 5-min washes of 50% ethanol before a final wash in 1X PBS for 10 min. Following the Thio-S stain, sections were treated with a standard indirect immunohistochemical protocol. Briefly, free-floating sections were immersed in normal blocking serum solution (5% normal goat serum with 0.2% TritonX-100 in 1X PBS) for 1 hr before an overnight incubation at 4°C in primary antibodies diluted in normal blocking serum solution.

Brain sections were stained with the following diluted primary antibodies against: ionized calcium-binding adapter molecule 1 (IBA1; 1:2000; 019-19741; Wako, Osaka, Japan), Aβ_1-16_(6E10; 1:2000; 8030001; BioLegend, San Diego, CA), glial fibrillary acidic protein (GFAP; 1:1000; AB134436; Abcam, Cambridge, MA), S100 β (1:200; AB41548; Abcam, Cambridge, MA), Fox 3 protein (NeuN; 1:1000; AB104225; Abcam, Cambridge, MA), Ctip2 (CTIP2; 1:300; AB18465; Abcam, Cambridge, MA), lysosome-associated membrane protein 1 (LAMP1; 1:200; AB25245; Abcam, Cambridge, CA), human tau (HT7; 1:1000; MN1000; Invitrogen, Waltham, MA), poly-tau (1:1000; Agilent Dako, Santa Clara, CA), phospho-tau Ser202, Thr 205 (AT8; 1:500; MN1020; Invitrogen, Waltham, MA), phospho-tau Thr217 (pTau217; 1:100; 44-744; Invitrogen, Waltham, MA), *Wisteria floribunda agglutinin* lectin (WFA; 1:1000; B-1355; Vector Labs, Burlingame, CA), parvalbumin (PV; 1:500; MAB1572; Millipore Sigma, Darmstadt, Germany).

Heat-induced antigen retrieval was necessary prior to staining sections with antibodies against Ctip2, AT8, and pT217. Sections were incubated in pre-heated citric acid buffer (pH 6.0) at 80°C for 30 min and allowed to cool to 20°C for 25 min. Afterwards, sections were rinsed in 1X PBS for 10 min and standard indirect immunohistochemical protocol was subsequently performed.

Amylo-Glo staining was used to confirm plaques with an additional marker. Free-floating sections were washed in 70% ethanol for 5 min and rinsed in deionized water for 2 min before being immersed for 10 min in Amylo-Glo RTD Amyloid Plaque Staining Reagent (1:100; TR-200-AG; Biosensis, Thebarton, South Australia) suspended in 0.9% saline solution. Afterwards, sections were washed in 0.9% saline solution for 5 minutes and briefly rinsed in deionized water for 15 sec before proceeding with standard indirect immunohistochemical protocol.

### Simplified Gallyas’ Method for Silver Staining

A simplified protocol of Gallyas’ method for silver staining was performed to stain neurofibrillary tangles (NFTs) in coronal brain sections of 3xTg-AD and wildtype mice at all timepoints (Kuninaka et al., 2015). A representative slice from each mouse was pooled together in wells containing slices from mice of the same experimental group (e.g. identical genotype, age, and sex) and were rinsed three times in 1X PBS for 5 min each. Following by a final wash of deionized water for 10 min, sections was submerged in 5% periodic acid for 3 min and then rinsed in deionized water for 5 min. The sections were incubated in silver iodide reagent (4% sodium hydroxide, 10% potassium iodide, 0.035% silver nitrate) chilled at 4°C for 1 min and subsequently washed twice in 0.5% acetic acid for 5 min each before rinsing in deionized water. A developer solution was prepared in advanced, combining a solution of 5% sodium carbonate and another solution of 0.19% ammonium nitrate, 0.2% silver nitrate, and 1% tungstosilicic acid, and 0.14% formaldehyde. The sections were incubated in this chilled developing solution for 15-25 min until they developed a pale brown or grey color, after which the development was stopped by immersion in 0.5% acetic acid for 5 min. Sections were mounted and dehydrated before being coverslipped using Cytoseal mounting medium. Microscope slides were allowed to dry before brightfield microscopy.

### Microscopy and Histological Analysis

Immunostained sections were mounted and coverslipped using Fluoromount-G either with or without DAPI (0100-20 or 0100-01, respectively; SouthernBioTech, Birmingham, AL). For whole-brain stitches, automated slide scanning via a Zeiss AxioScan.Z1 was used with a 10X, 0.45 NA objective. Scanned images were corrected for shading, stitched together, and exported using ZEN 2.3 slidescan software for representative images. High resolution confocal images of immunofluorescence images were also captured using a Leica TCS SPE-II confocal microscope with a 10X, 0.3 NA objective lens and LAS-X software. Max projections of 20X Z-stacks of subiculum and visual cortex regions were imaged per section per mouse and used for both Bitplane Imaris quantification and representative images. High resolution images of sections stained using the simplified Gallyas method were imaged using a Zeiss Axioscope 5 with a 20X objective. Images were exported using AxioVision LE64 and used for both FIJI ImageJ quantification and representative images.

#### Imaris Quantitative Analysis

Confocal images of each brain region were quantified automatically using the spots module within the Imaris v9.7 software (Biplane Inc. Zürich, Switzerland) then normalized to the area of the field-of-view (FOV). Amyloid burden was assessed by measuring both the total Thio-S^+^ plaque number normalized to FOV area and their volume via the surfaces module in Imaris software. Similarly, volumetric measurements (i.e. Thio-S^+^ plaque volume, PNN volume, etc.) were also acquired automatically utilizing the surfaces module on confocal images of each brain region. Quantitative comparisons between experimental groups were carried out in sections stained simultaneously.

#### FIJI ImageJ Analysis

Brightfield microscopy images of silver-stained brain sections were first opened in ImageJ software (Schneider et al., 2012) and converted to 16-bit grey-scale. The threshold feature was adjusted and used to distinguish cells from background, which were analyzed using the analyze particles feature. As a result, the ROI manager includes a summary box containing the total number of cells per image.

### Biochemical Analysis

Micro-dissected hippocampal and cortical regions of each mouse were flash-frozen and processed for biochemical analysis. Samples were pulverized using a Bessman Tissue Pulverizer kit. Pulverized hippocampal tissue separated for biochemical analysis was homogenized in 150µL of Tissue Protein Extraction Reagent (TPER; Life Technologies, Grand Island, NY), while cortical tissue was homogenized in 1000µL/150 mg of TPER. Together with protease (Roche, Indianapolis, IN) and phosphatase inhibitors (Sigma-Aldrich, St. Louis, MO), the homogenized samples were centrifuged at 100,000 g for 1 hr at 4°C to generate TPER-soluble fractions. For formic acid-fractions, pellets from TPER-soluble fractions were homogenized in 70% formic acid: 75µL for hippocampal tissue or half of used TPER volume for cortical tissue. Afterwards, samples were centrifuged again at 100,000 g for 1 hr at 4°C. Protein concentration in each soluble or insoluble fraction was determined via Bradford Protein Assay (Bradford, 1976); (Green et al., 2011).

#### Electrochemiluminescence-linked immunoassay

Quantitative biochemical analyses of human Aβ soluble and insoluble fraction levels were acquired using the V-PLEX Aβ Peptide Panel 1 (6E10) (K15200G-1; Meso Scale Discovery, Rockville, MD). Plates were prepared and read according to the manufacturer’s instructions.

### Statistics

Every reported *n* represents the number of independent biological replicates. The sample sizes are similar with those found in prior studies conducted by MODEL-AD (Forner et al., 2021) and were not predetermined using statistical methods. Behavioral, immunohistochemical, and biochemical data were analyzed using Student’s t-test, one-way ANOVA, or two-way ANOVA via Prism v.9 (GraphPad, La Jolla, CA). Bonferonni-Šídák and Tukey’s post hoc tests were utilized to examine biologically relevant interactions from the two-way ANOVA. * p ≤ 0.05, * * p ≤ 0.01, * * * p ≤ 0.001, * * * * p ≤ 0.0001. Statistical trends are accepted at p < 0.10 (^#^). Data are presented as raw means and standard error of the mean (SEM).

### Data Records

The protocols, data, and results are available via the AD Knowledge Portal (https://adknowledgeportal.synapse.org). The AD Knowledge Portal is a platform for accessing data, analyses, and tools generated by the Accelerating Medicines Partnership (AMP-AD) Target Discovery Program and other National Institute on Aging (NIA)-supported programs to enable open-science practices and accelerate translational learning. The data, analyses and tools are shared early in the research cycle without a publication embargo on secondary use. Data is available for general research use according to the following requirements for data access and data attribution (https://adknowledgeportal.org/DataAccess/Instructions).

Data can be accessed in an interactive matter at UCI Mouse Mind Explorer (admodelexplorer.org).

## Results

### 3xTg-AD mice display minimal behavioral deficits

3xTg-AD and WT control mice were aged to 4, 12, and 18-months of age and subjected to a battery of tasks to test for cognitive and behavioral differences. Further characterization of the 3xTg-AD mouse model was obtained through a systemic pipeline of analyses investigating differences in long-term potentiation (LTP), immunohistochemistry, biochemistry, and gene expression.

No differences are found between 3xTg-AD mice and age-matched WT negative control mice tasked in the elevated plus maze (EPM), a task in which the 5xFAD mice show profound deficits from 4 mo of age (Forner et al. 2021). At all timepoints, 3xTg-AD and WT mice exhibit similar time spent in open and closed arms (Fig. 1A-B), although both genotypes spend the expected increased time in closed arms rather than open arms. No sex differences are found in 3xTg-AD mice nor age-matched controls in the EPM task. No changes are observed between 3xTg-AD and control mice in Y maze tasks, which tests working memory, as both groups enacted similar numbers of alterations at all timepoints (Fig. 1C). No sex differences are observed during these Y maze tasks either. However, motor differences in 18-month-old 3xTg-AD mice are detected during the open field tasks as an increased distance traveled and increased velocity compared to age-matched WT controls (Fig. 1D-E). Additionally, no differences are found between 3xTg-AD and WT control mice performing contextual fear conditioning tasks at all timepoints nor are sex differences observed in these tasks (Fig. 1G).

**Figure 1.**
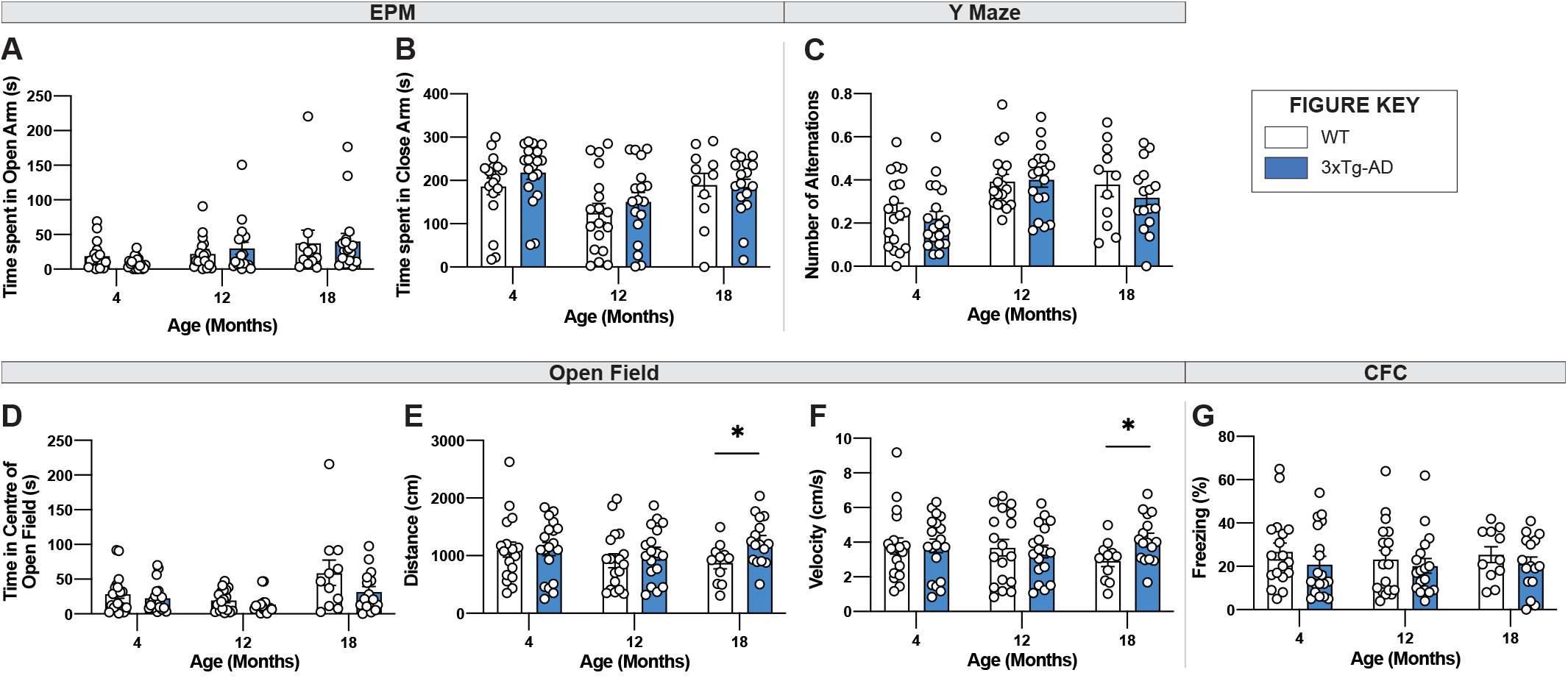
Behavioral tasks reveal age related changes in 18-month-old 3xTg-AD mice. **(A-B)** Elevated plus maze tasks on either WT or 3xTg-AD mice reveal no age-, genotype-, or sex-related differences in time spent in the open arms **(A)** or closed arms **(B). (C)** Y maze tasks on either WT or 3xTg-AD mice reveal age-related effects on the number of alterations, but no differences between genotypes or sex. (**D-F**) Open field testing reveals genotype-related increase in distance traveled (**E**) and velocity (**F**) in 18-month-old 3xTgAD mice. However, no sex differences within genotypes were observed. **(G)** There is no effect of either age, sex, or genotype on the contextual fear conditioning task. Data are represented as mean ± SEM. * p ≤ 0.05.

### 3xTg-AD mice demonstrate age-associated LTP impairments

We examined long-term potentiation in acute hippocampal slices from 4-, 12-, and 18-month-old male and female 3xTg-AD and WT mice. As early as 4 months of age, there is a marked decrease in LTP 50-60 min post theta burst stimulation (TBS) in stratum radiation of area CA1 in slices from 3xTg-AD mice relative to WT controls (Fig 2A-D). This overall group effect is largely driven by female mice (Fig. 2E, H). At 12 months of age, the variability in LTP at 1 hr post-TBS is great, however, a deficit in LTP is still present in females, but not males (Fig. 2F, H). By 18 months of age, LTP in slices from both male and female 3xTg-AD is-significantly reduced as compared to age-matched control WT slices (Fig. 2G, H). Analysis of the mean potentiation during the last 10 min of recording following TBS revealed that there are significant group differences at 4 and 18 months of age (Fig. 2D). Moreover, the level of potentiation in slices from 18-month-old 3xTg-AD mice is significantly reduced as compared to slices from 4- and 12- month-old 3xTg-AD.

**Figure 2.**
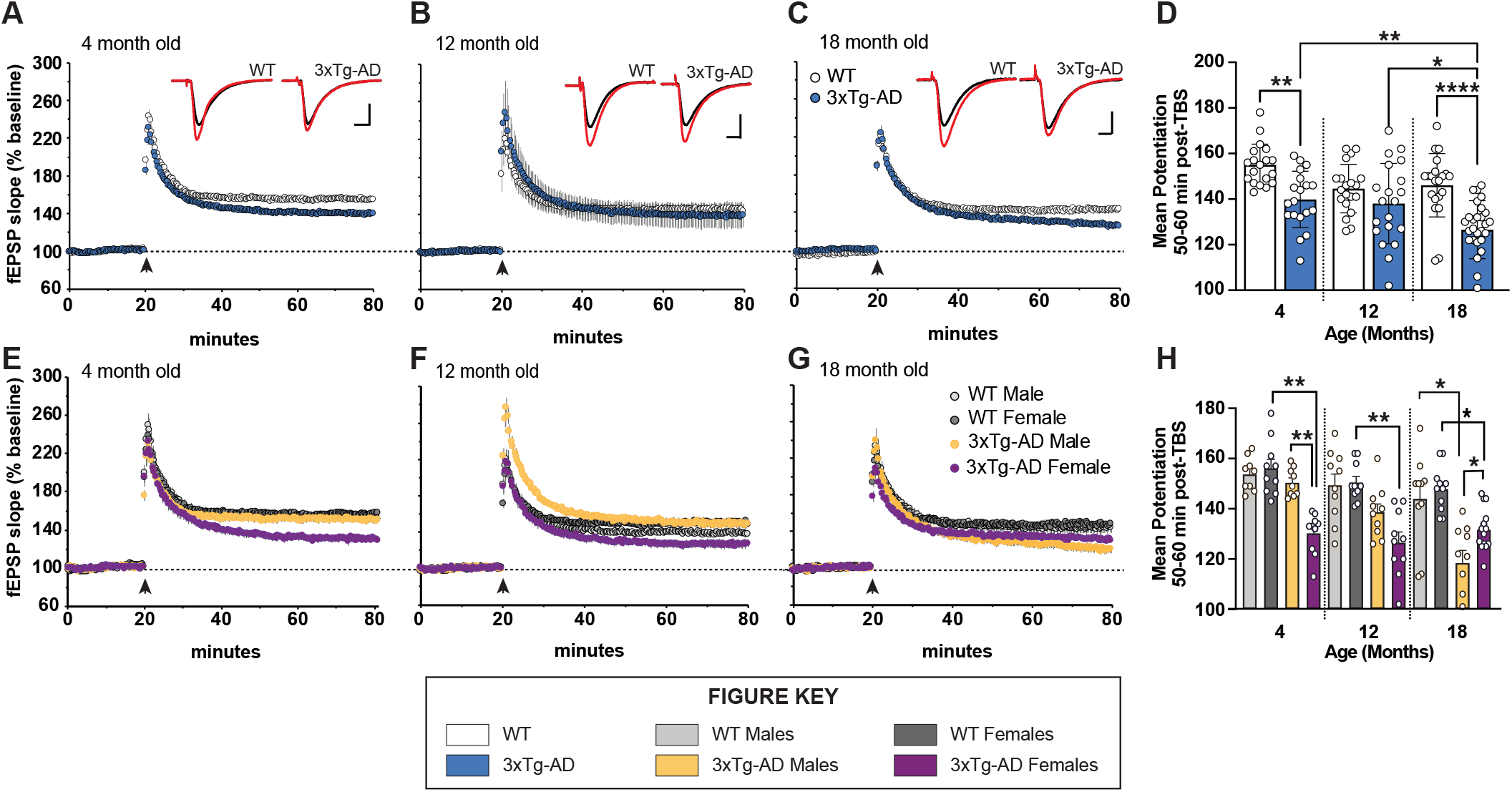
LTP Impairments in 3xTg-AD mice at ages 4,12, and 18 months in a sex-dependent manner. **(A-C, E-G)** Hippocampal slices were collected from 4-, 12-, and 18-month old (mo) male and female WT and 3xTg-AD mice and were used to measure long-term potentiation (LTP) in the stratum radiatum in area CA1. Following a 20 min stable recording of field excitatory postsynaptic potentials (fEPSP), LTP was induced by applying 5 theta bursts (black arrow: each burst containing 4 100Hz pulses with each burst separated by 200 ms) and recording of baseline stimulation was resumed for an additional 60 min. **(D, H)** fEPSP potentiation averaged during the last 10 min of recording in slices from male and female WT and 3xTg-AD mice at ages 4, 12, and 18 mo. * p<0.05, * * p<0.005, * * * * p<0.0001.

### Age-associated increases in fibrillar Aβ plaque burden and size

Coronal brain sections from 3xTg-AD and control mice of both sexes at 4-, 12-, and 18-month timepoints were stained with Thio-S for the characterization of fibrillar Aβ plaques (Fig. 3A-B). As expected, no plaques are detected in WT control mice. Thio-S plaques are only seen in a subset of female 3xTg-AD hippocampi at 12 months of age and most females by 18 months (Fig. 3C-D). Male mice have far fewer plaques, with only a small number of males showing Thio-S^+^ plaques at 18 months. Notably, plaques are not detected outside of the subiculum in any animal (Fig. 3A). Consistent with these observations, the total volume of Thio-S+ plaques is significantly increased in 18-month-old 3xTg-AD mice compared to other timepoints, an effect largely driven by female 3xTg-AD mice (Fig. 3E-F). A similar sex-specific effect is seen with the average volume per Thio-S^+^ plaque (Fig. 3G-H), consistent with an increase in plaque size between 12-month and 18-month timepoints.

**Figure 3.**
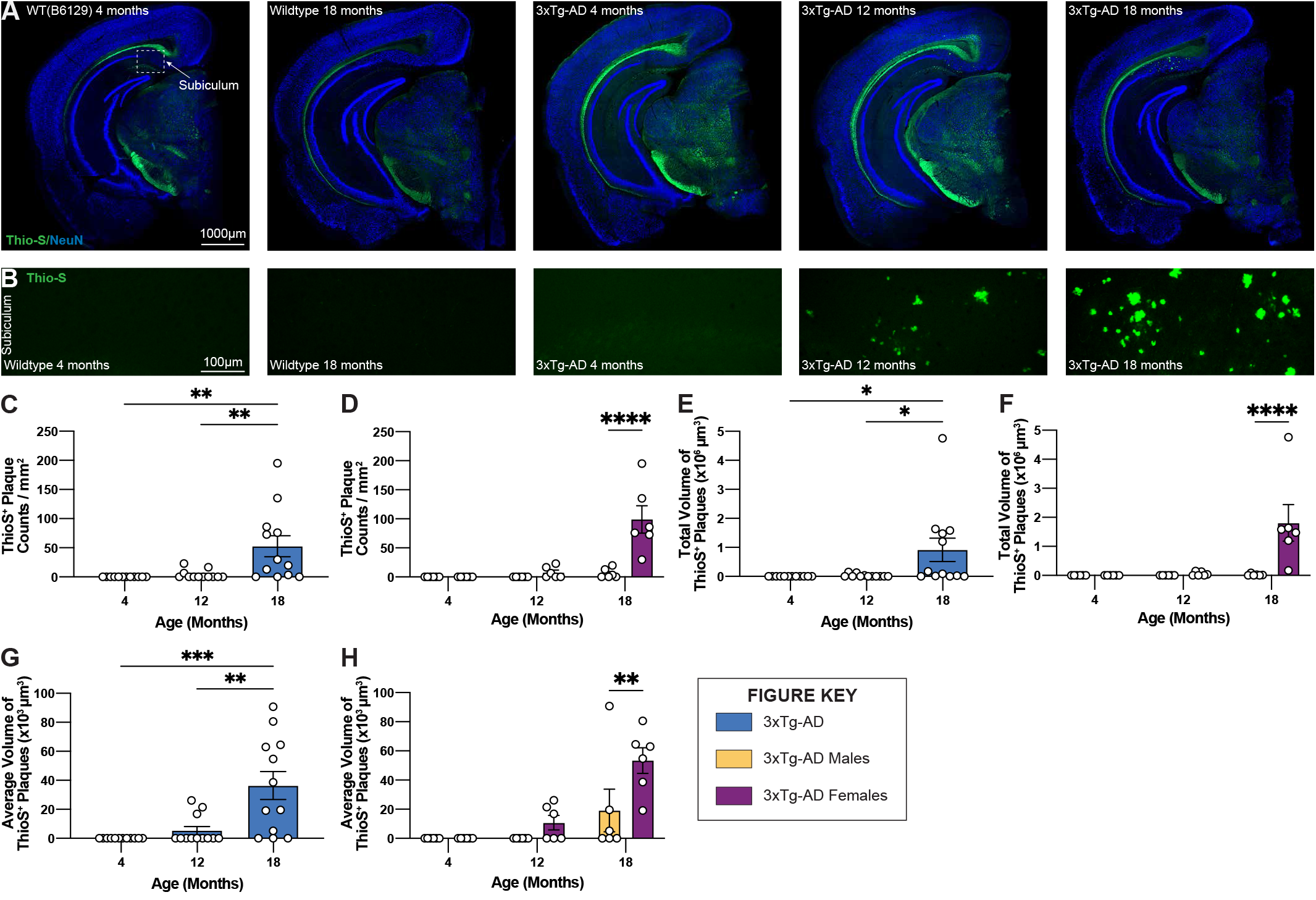
Fibrillar amyloid plaques increase in size and number in 18-month-old female 3xTg-AD mice. 3xTg-AD plaque burden was assessed with Thioflavin-S staining at each time point. **(A)** Representative stitched brain hemispheres of WT and 3xTg-AD mice shown with Thio-S^+^ staining at 4- and 18-month, and 4, 12, and 18-month timepoints, respectively counterstained with NeuN. **(B)** Representative confocal images of Thio-S plaques in subiculum hippocampal regions of WT and 3xTg-AD mice across timepoints displaying increased number of fibrillar amyloid plaques. **(C-D)** Quantification for density of Thio-S^+^ plaques in the subiculum hippocampal region per square millimeter by genotype and sex. **(E-F**) Quantification of total volume of Thio-S^+^ plaques by genotype and sex demonstrating age- and genotype-associated increases in total volume of plaques. (**G-H**) Quantification of average volume of Thio-S^+^ plaques showing an increase in the average volume of a plaque related to age and genotype. n = 6 mice per genotype/age/sex. Data are represented as mean ± SEM. * p ≤ 0.05, * * p ≤ 0.01, * * * p ≤ 0.001, * * * * p ≤ 0.0001.

Aβ40 and Aβ42 were quantified in detergent soluble and insoluble fractions of micro-dissected hippocampus and cortical tissue. The soluble and insoluble forms correspond to recently produced Aβ, and Aβ contained within plaques, respectively. As anticipated, a substantial increase in both Aβ40 and Aβ42 is found in soluble fractions of both hippocampus and cortex in 18-month 3xTg-AD mice (Fig. 4A-H). Consistent with plaque load, 18-month-old female 3xTg-AD mice display elevated soluble Aβ40 and Aβ42 in hippocampus compared to 3xTg-AD male mice (Fig. 4B, F). Soluble Aβ40, but not Aβ42, is slightly, but significantly increased in the cortices of female 3xTg-AD mice compared to male 3xTg-AD mice (Fig. 4D, H).

**Figure 4.**
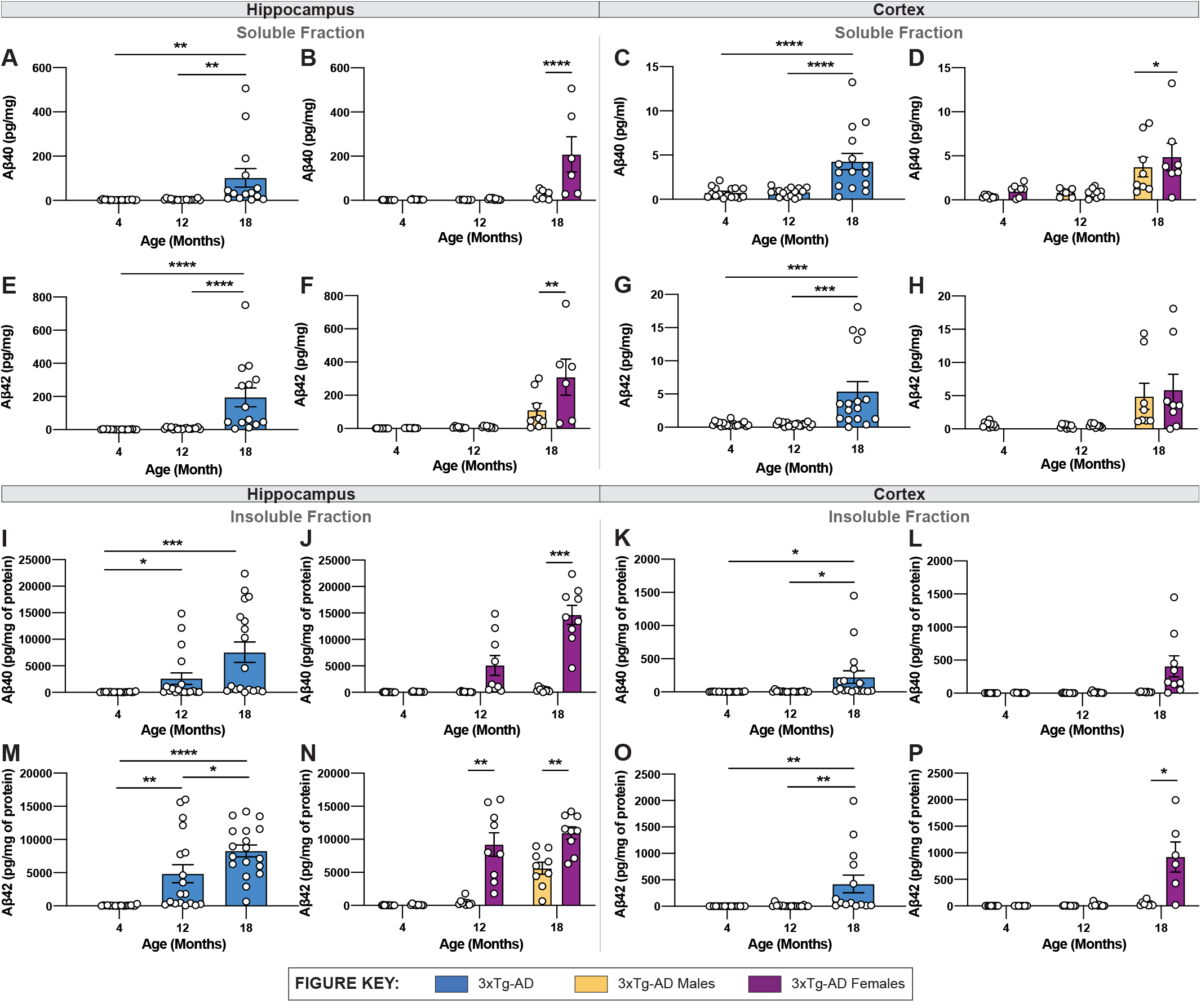
Quantification of Aβ isoforms in 3xTg-AD mice of different age and sex. Aβ was quantified in micro-dissected hippocampi and cortices via Mesoscale Multiplex technology. **(A-H)** Aβ40 and Aβ42 were measured in the soluble fraction of hippocampus (**A-B, E-F**) and cortex **(C-D, G-H**), respectively, with age-related increases in Aβ40 and Aβ42 shown in hippocampus and cortex of 3xTg-AD mice between sexes. **(I-P)** An age-related increase in insoluble Aβ40 and Aβ42 was also observed in hippocampus **(I-J, M-N**) and cortex **(K-L, O-P)** of 3xTg-AD mice between sexes. Data are represented as mean ± SEM. * p ≤ 0.05, * * p ≤ 0.01, * * * p ≤ 0.001, * * * * p ≤ 0.0001.

In tandem with increased plaque density, insoluble Aβ40 and Aβ42 are also increased in hippocampal and cortical regions in 3xTg-AD mice with age (Fig. 4I-P). Insoluble Aβ40 and Aβ42 progressively increase between 4-, 12- and 18-month-old 3xTg-AD mice (Fig. 4i, m). As with soluble Aβ, there is a clear sex difference in the quantity of insoluble Aβ, with 18-month-old female 3xTg-AD mice showing significantly more Aβ40 and Aβ42 in hippocampal tissue (Fig. 4J, N).

### Accumulation of human tau and poly-tau in hippocampal regions

Brain sections from 3xTg-AD mice at 4-, 12-, and 18-month timepoints were stained for human tau using HT7 antibody and combined human & mouse tau protein using poly-tau antibody. As expected, in wildtype mice expression of human tau or somatodendritic accumulation of murine tau is not observed (Fig. 5A-B). In contrast, 3xTg-AD mice, that overexpress mutant human tau (P301L), display substantial staining using HT7^+^ and poly-tau^+^ antibodies in the somatodendritic compartment of hippocampal CA1 region neurons (Fig. 5A-B). Quantification of the total volume of HT7^+^ cells demonstrates age-related differences in 3xTg-AD mice, increased at 12-months but decreased at 18-months (Fig. 5C). This effect is mostly contributed by female 3xTg-AD mice, which have greater total volumes of HT7^+^ neurons compared to male 3xTg-AD mice at 4- and 12-month timepoints. Interestingly, this sex difference diminishes at 18-months of age (Fig. 5D). Unlike total volumes of HT7^+^ cells, the total volume of cells staining for poly-tau antibody display an age-associated increase in hippocampal CA1 regions of 18-month 3xTg-AD mice (Fig. 5E). Once again, sex differences are evident at 4- and 12-month timepoints as female 3xTg-AD mice exhibit greater total volumes compared to males, an effect that diminishes at the 18-month timepoint (Fig. 5F).

**Figure 5.**
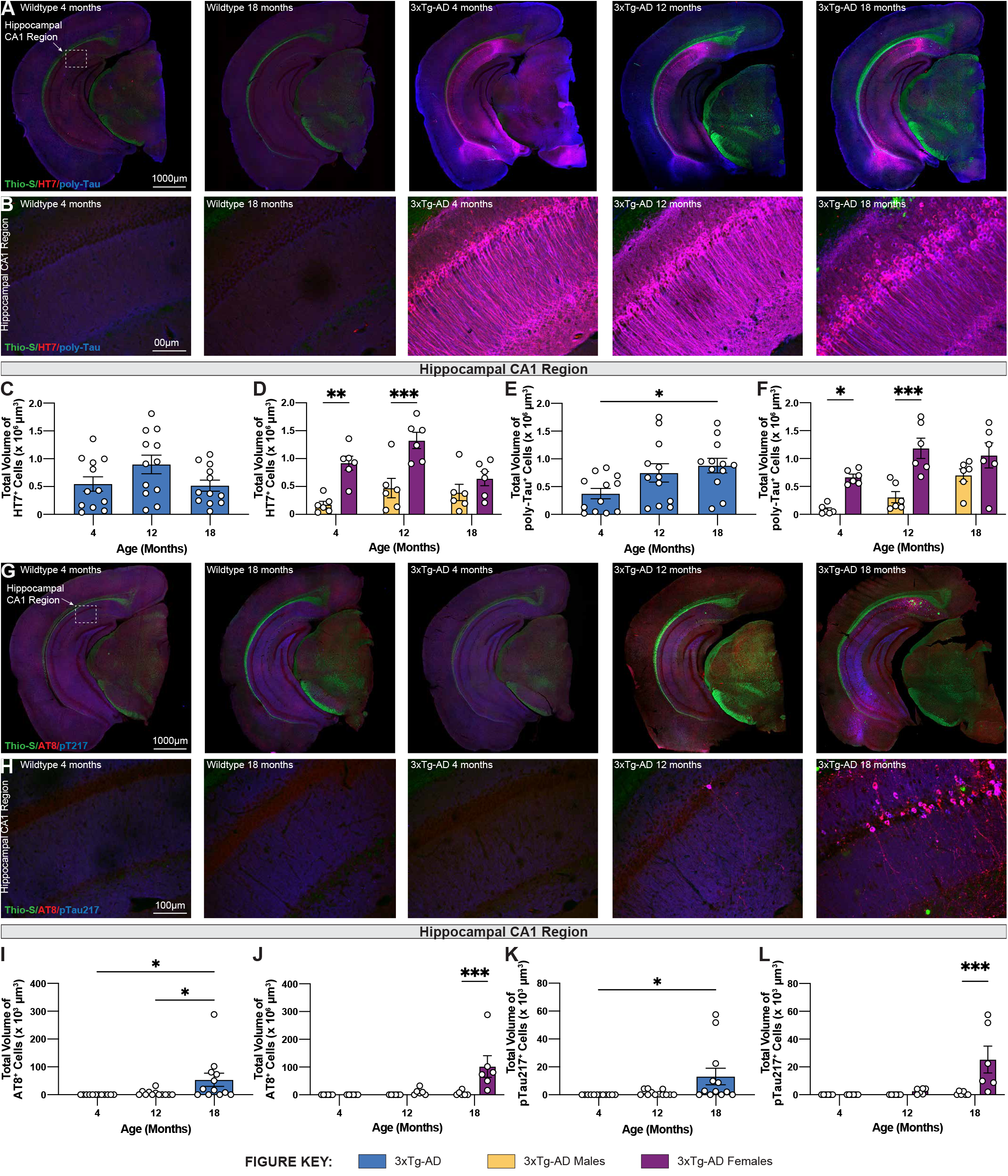
Phosphorylated tau increases in volume in 18-month-old female 3xTg-AD mice. Accumulation of human tau was assessed by immunostaining using HT7 and poly-tau antibody, while phosphorylated tau was assessed using AT8 and phospho-Tau Thr 217 antibody, respectively. **(A)** Representative stitched brain hemispheres of WT and 3xTg-AD mice stained with Thio-S/HT7/poly-Tau at 4- and 18-month and 4, 12, and 18-month timepoints, respectively. **(B)** Representative confocal images of HT7^+^ and poly-Tau^+^ cells in the hippocampal CA1 regions of WT and 3xTg-AD mice across respective timepoints. **(C-D)** HT7 immunostaining reveals age-related changes in total volume of HT7^+^ cells in 3xTgAD mice between sexes. **(E-F)** Poly-Tau immunostaining reveals age-related changes in total volume of poly-Tau^+^ cells in 3xTg-AD mice between sexes. **(G)** Representative stitched brain hemispheres of WT and 3xTg-AD mice stained with Thio-S/AT8/pT217 at 4- and 18-month and 4, 12, 18-month timepoints, respectively. **(H)** Representative confocal images of AT8^+^ and Thr217^+^ cells in the hippocampal CA1 regions of WT and 3xTg-AD mice across respective timepoints. **(I-J)** AT8 immunostaining reveals age-related changes in total volume of AT8^+^ cells in 3xTg-AD mice between sexes. **(K-L)** Thr217^+^ immunostaining reveals age-related changes in total volume of Thr217 cells in 3xTg-AD mice between sexes. n = 5-6 mice per genotype/age/sex. Data are represented as mean ± SEM. * p ≤ 0.05, * * * p≤0.001.

### Age-associated accumulation of phosphorylated tau species in hippocampal regions

To characterize hyperphosphorylated tau in 3xTg-AD mice, brain sections from each 3xTg-AD mouse at 4-, 12-, and 18-month timepoints were stained for phosphorylated tau species using antibodies against paired helical filament tau phosphorylated at serine 202 and threonine 205 (via antibody AT8). Phospho-tau Thr217 (pTau217) was recently found to be present in the blood plasma of Alzheimer’s Disease patients (Palmqvist et al., 2020); (Barthelemy et al., 2020). Therefore, an antibody against pTau217 was also used to detect this post-translational modification in the brain sections of 3xTg-AD mice at each timepoint. No AT8^+^ and pTau217^+^ cells are observed in the brains of age-matched wildtype mice. In contrast, a strong signal to phosphorylated tau was present in the brains of 18-month-old 3xTg-AD mice (Fig. 5G-H). Quantification of the total volume of AT8^+^ and pTau217^+^ cells show significant age-related accumulation of phosphorylated tau species in the hippocampal CA1 region of 3xTg-AD mice (Fig. 5I, K). Again, accumulation of phospho-tau species is driven largely by female 3xTg-AD mice (Fig. 5J, L)

### Development of neurofibrillary tangles in the CA1 of aged female 3xTg-AD mice

To visualize neurofibrillary tangles (NFTs), brain sections from each 3xTg-AD mouse at 4-, 12-, and 18-month timepoints were stained using a simplified Gallyas’ silver stain. As expected, no NFTs are detected in age-matched wildtype controls but are present in female 3xTg-AD hippocampi at 18 months of age (Fig. 6a-d), but not in male mice, or in other brain regions (Fig. 6A-D).

**Figure 6.**
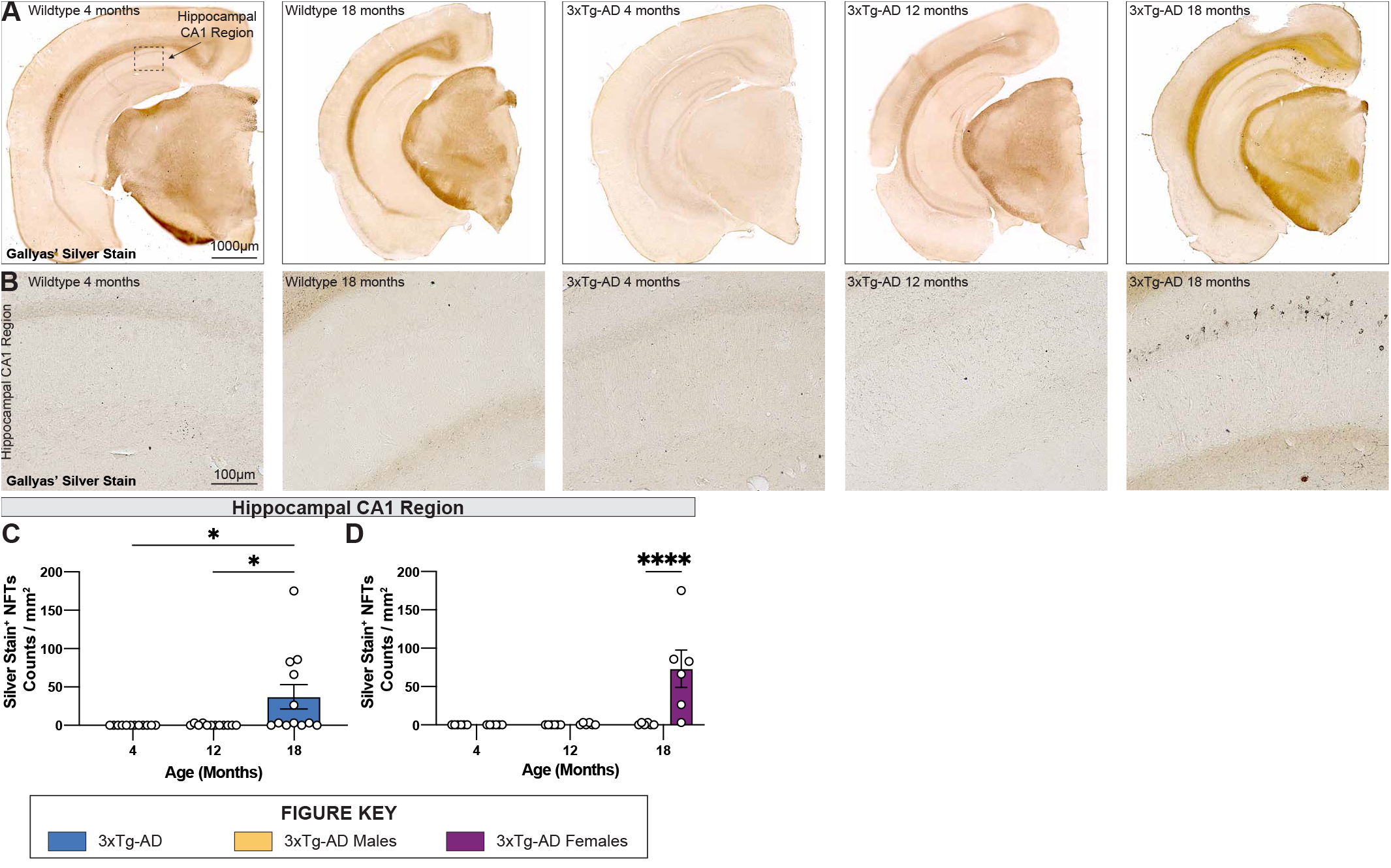
Neurofibrillary tangles increase in number in 18-month-old female 3xTg-AD mice. Accumulation of neurofibrillary tangles (NFTs) was assessed using Gallyas’ silver staining method at each timepoint. **(A)** Representative stitched brain hemispheres of WT and 3xTg-AD mice at 4- and 18-month, and 4, 12, and 18-month timepoints, respectively. **(B)** Representative brightfield images of silver stained NFTs in hippocampal CA1 regions of WT and 3xTg-AD mice across each timepoint showing progressive accumulation of NFTs. **(C-D)** Quantification for density of silver stained NFTs in hippocampal CA1 regions per square millimeter by genotype and sex. n = 6 mice per genotype/age/sex. Data are represented as mean ± SEM. * p ≤ 0.05, * * * * p ≤ 0.0001.

### Age-related gliosis in 3xTg-AD mice

To investigate changes in glial cells in the brains of 3xTg-AD mice, sections of wildtype and 3xTg-AD mice at all timepoints were immuno-stained using antibodies against the microglial marker IBA1 (Fig. 7A), or astrocytic markers GFAP and S100β (Fig. 7G). An increase in microglial density in the subiculum accompanied the presence of dense core plaques in 18-month-old female 3xTg-AD mice (Fig. 7B-D). Plaques are not observed in the cortex of 3xTg-AD mice at any age (up to 18 mo of age), and no increase in microglial density was seen within cortical regions. An age-associated reduction in microglial density is observed in 12 and 18-month old WT females compared to male mice (Fig. 7E-F). In this study, eighteen-month-old male 3xTg-AD mice also have reduced densities of microglia in the visual cortex compared to WT controls (Fig. 7F).

**Figure 7.**
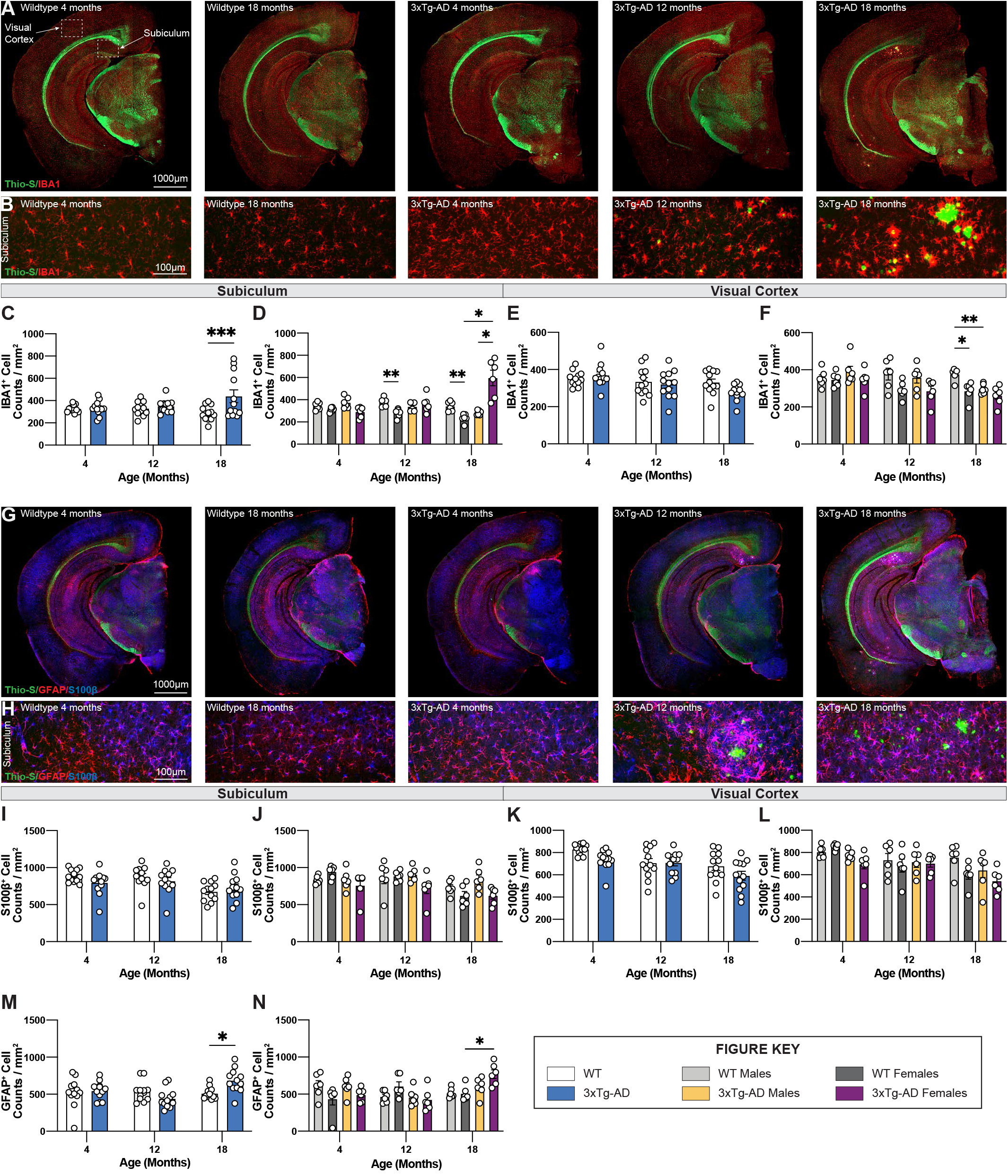
Immunostaining of microglia and astrocytes. Brains of mice at each timepoint were coronally sectioned and immunostained for IBA1, GFAP, and S100β to identify changes in microglia or astrocytes. **(A)** Representative stitched images of brain hemispheres of WT (4-, 18-month) and 3xTg-AD mice (4, 12, 18-month) stained with Thio-S/IBA1. **(B)** Representative images of IBA1^+^ cells surrounding Thio-S^+^ in subiculum hippocampal regions of WT and 3xTg-AD mice at indicated timepoints. **(C-F)** Age-related changes microglial density in both WT and 3xTg-AD, and differences between genotypes in subiculum hippocampal and cortical regions. **(G)** Representative images of brain hemispheres of WT (4-, 18-month) and 3xTg-AD mice (4, 12, 18-month) stained with Thio-S/GFAP/S100β. **(H)** Representative images of GFAP^+^ and S100β^+^ cells surrounding Thio-S^+^ plaques in subiculum hippocampal regions of WT and 3xTg-AD mice at indicated timepoints. **(I-N)** Astrocyte density as assessed via S100β **(I-L)** and GFAP **(M-N)** staining in the subiculum hippocampal and cortical regions. Age-related changes in both WT and 3xTg-AD astrocytic density, and differences between genotypes in both subiculum hippocampal and cortical regions. n = 5-6 mice per genotype/age/sex. Data are represented as mean ± SEM. * p ≤ 0.05, * * p ≤ 0.01, * * * p ≤ 0.001.

Densities and distribution of homeostatic (S100β +ve) and reactive (GFAP +ve) astrocytes were also measured by immunostaining. S100β is a transcription factor localized in the nucleus of all astrocytes while GFAP is expressed in both hippocampal astrocytes and “reactive” astrocytes in the cortex (Fig. 7G). Within the subiculum, homeostatic astrocytes (S100β +ve) populations are unchanged across all timepoints whereas reactive (GFAP +ve) astrocytes accumulate around Thio-S^+^ plaques (Fig. 7H). The density of S100β^+^ astrocytes in the subiculum hippocampal region across each timepoint was stable (Fig. 7 I-J) whereas the density of GFAP+ve astrocytes increases at the 18-month timepoint (Fig. 7 M-N). Interestingly, S100β^+^ astrocyte densities decrease with age in cortical regions of both 3xTg-AD and wildtype mice with a general trend particularly in both 3xTg-AD and WT female mice at 18 months (Fig. 7K-L). As expected, no reactive astrocytes are detected in cortical regions due to the lack of Thio-S+ plaques; therefore, no quantification was performed for GFAP immunostaining in cortical regions.

### Age-dependent change in perineuronal net volume and loss of PV^+^ interneurons in 3xTg-AD brains

We previously reported plaque-induced and microglia-mediated loss of perineuronal nets (PNNs) in the 5xFAD mouse model of AD as well as in human AD prefrontal cortex (Crapser et al., 2020). Age-associated changes in PNNs were investigated in hippocampal and cortical regions of 3xTg-AD and wildtype mice at each timepoint using the lectin Wisteria floribunda agglutinin (WFA), as well as parvalbumin (PV) which stains PV+ interneurons, LAMP1 – a marker of dystrophic neurites, and amyloglo for dense core plaques (Fig. 8A-B). In accordance, 18-month-old 3xTg-AD mice show significant reductions in PNN volume in the subiculum driven mainly by female mice (Fig. 8C-D) coincident with the appearance of plaques, while no changes are seen in the visual cortex (Fig. 8E-F) which lack plaques. Interestingly, in WT mice females maintain PNN volumes across their lifespans in the subiculum, while male mice show age-dependent reductions (Fig. 8C-D).

**Figure 8.**
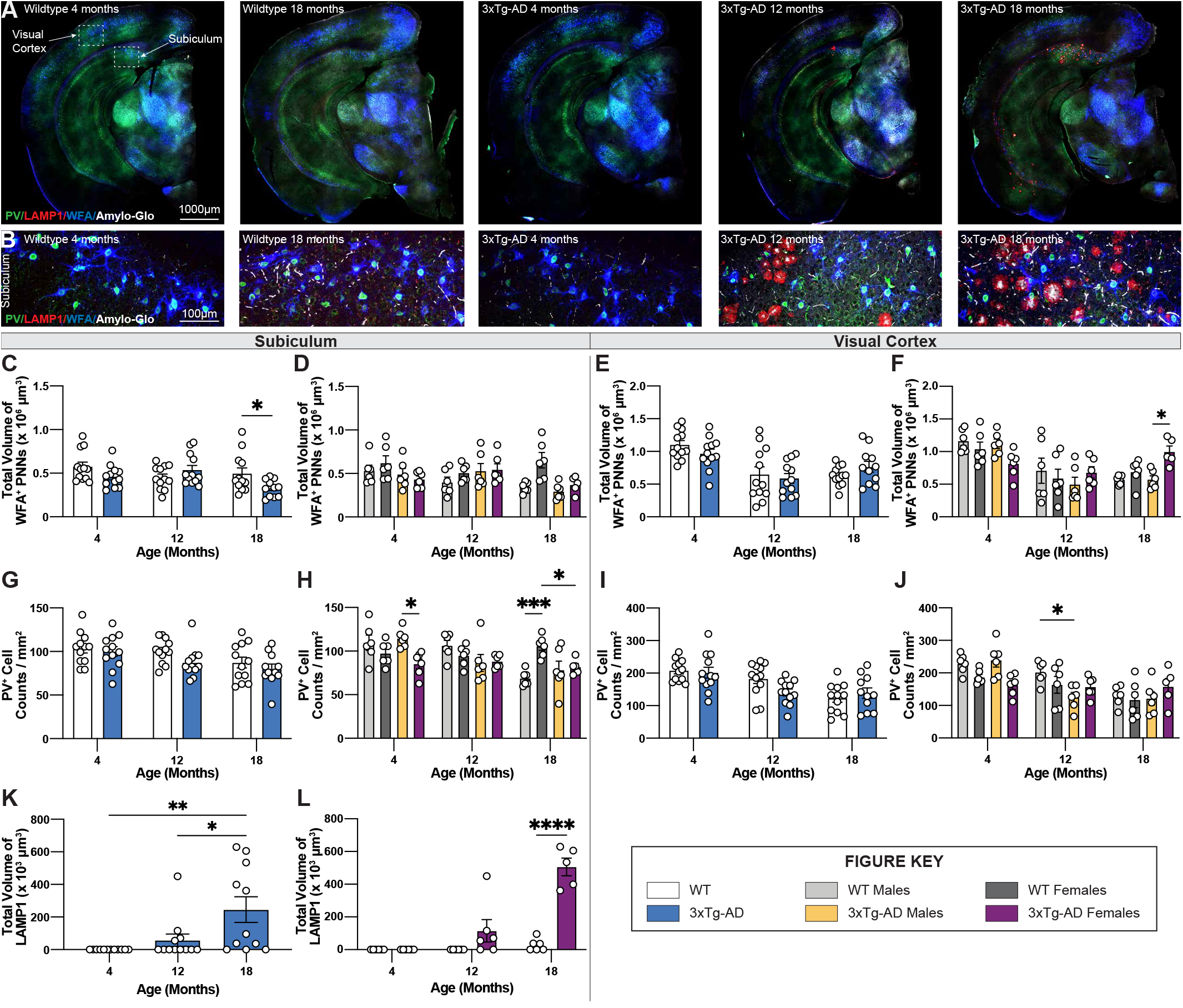
PNN and LAMP1. Perineuronal nets (PNNs) and parvalbumin (PV) interneurons were assessed with immunostaining using WFA and PV antibody, while lysosomes were assessed with immunostaining using LAMP1 antibody. **(A)** Representative images of WT (4-, 18-month) and 3xTg-AD mice (4, 12, 18-month) stained with PV/LAMP1/WFA/Amylo-Glo. **(B)** Representative images of WFA^+^ PNNs surrounding PV^+^ neurons around LAMP1^+^ lysosomes in subiculum hippocampal regions of WT and 3xTg-AD mice across respective timepoints. **(C-F)** Age-related change in total volume of WFA+ PNNs in 3xTg-AD mice. **(G-J)** Immunostaining for PV+ neurons demonstrate age-related changes in the density of PV+ neurons in both WT and 3xTg-AD mice, and differences between genotypes in subiculum and cortical regions. n = 5-6 mice per genotype/age/sex. **(K-L)** LAMP1 immunostaining reveals age- and sex-associated increases in the total volume of LAMP1+ lysosomes in the subiculum brain region of 3xTg-AD mice. Data are represented as mean ± SEM. * p ≤ 0.05, * * p ≤ 0.01, * * * p ≤ 0.001, * * * * p ≤ 0.0001.

In 5xFAD mice, loss of PNNs preceded loss of parvalbumin^+^ (PV^+^) interneurons (Crapser et al., 2020). Likewise, significant reductions in PV+ interneurons are seen in the subiculum of 18-month-old female 3xTg-AD compared to WT mice (Fig. 8G-H). Notably, mirroring the age-related loss of WFA+ PNN’s in male WT mice is an age-related loss of PV+ neurons, which are maintained in female mice (Fig. 8gG-H). Age related reductions in PV+ interneurons in the visual cortex are seen in both male and female mice, but no additional reductions in 3xTg-AD mice (Fig. 8I-J), presumably due to the lack of plaque pathology in that region.

### Age-dependent accumulation of dystrophic neurites in 3xTg-AD hippocampus

Dystrophic neurites surround dense core plaques, which can be visualized through immunostaining with LAMP1. Immunofluorescence staining was performed using an antibody against LAMP1 and Amylo-Glo, a fluorescent tracer for Aβ plaques, on brain sections of 3xTg-AD and wildtype mice at 4-, 12-, and 18-month timepoints (Fig. 8A-B). Neither dystrophic LAMP1^+^ dystrophic neurites nor Amylo-Glo^+^ plaques accumulate in wildtype mice. In contrast, 3xTg-AD mice show the presence of dystrophic neurites alongside plaques within subiculum brain regions (Fig. 8B). Quantifying the total volumes of LAMP1 staining reveals significant age-associated increase in dystrophic neurites in 3xTg-AD mice, which is mostly due to female 3xTg-AD mice (Fig. 8K-L). As expected, this pattern mirrors the age-associated increase of Thio-S^+^ dense core plaques in female 3xTg-AD mice.

### Changes in gene expression found in 3xTg-AD brains

Bulk tissue gene expression was measured via RNA-seq, from microdissected hippocampi, and can be explored in an interactive and searchable fashion at https://admodelexplorer.synapse.org. Comparisons between bulk tissue and single cell/single nucleus RNA-seq from 3xTg-AD brains are explored extensively in a companion publication (Balderrama-Gutierrez et al., 2021).

### Comparison of 3xTg-AD and 5xFAD mouse models

To enable a comparison of an onset of Alzheimer’s-related histopathology between two established animal models of AD, representative sections of 3xTg-AD and 5xFAD mice with their respective wildtype B6/129 and B6J controls were immuno-stained for Thio-S^+^ dense core plaques and IBA1^+^ microglia. Representative images of brains of each wildtype mouse display minimal differences between B6/129 and B6J controls. Abundant accumulation of Thio-S^+^ dense core plaques is apparent in hemizygous 5xFAD mice at 4-months of age compared to 3xTg-AD mice at 18-months of age (Fig. 9A-B). Analysis of subiculum hippocampal regions reveals an increase in the density of Thio-S^+^ plaques in hemizygous 5xFAD mice compared to 3xTg-AD (Fig. 9C-D). These differences are also observed in the microgliosis of IBA1^+^ cells surrounding dense core Thio-S^+^ plaques. As expected, a significant increase in IBA1^+^ microglia density accompanies the presence of Thio-S^+^ plaques in hemizygous 5xFAD mice as early as 4-months of age versus 18-month-old 3xTg-AD mice (Fig. 9E).

**Figure 9.**
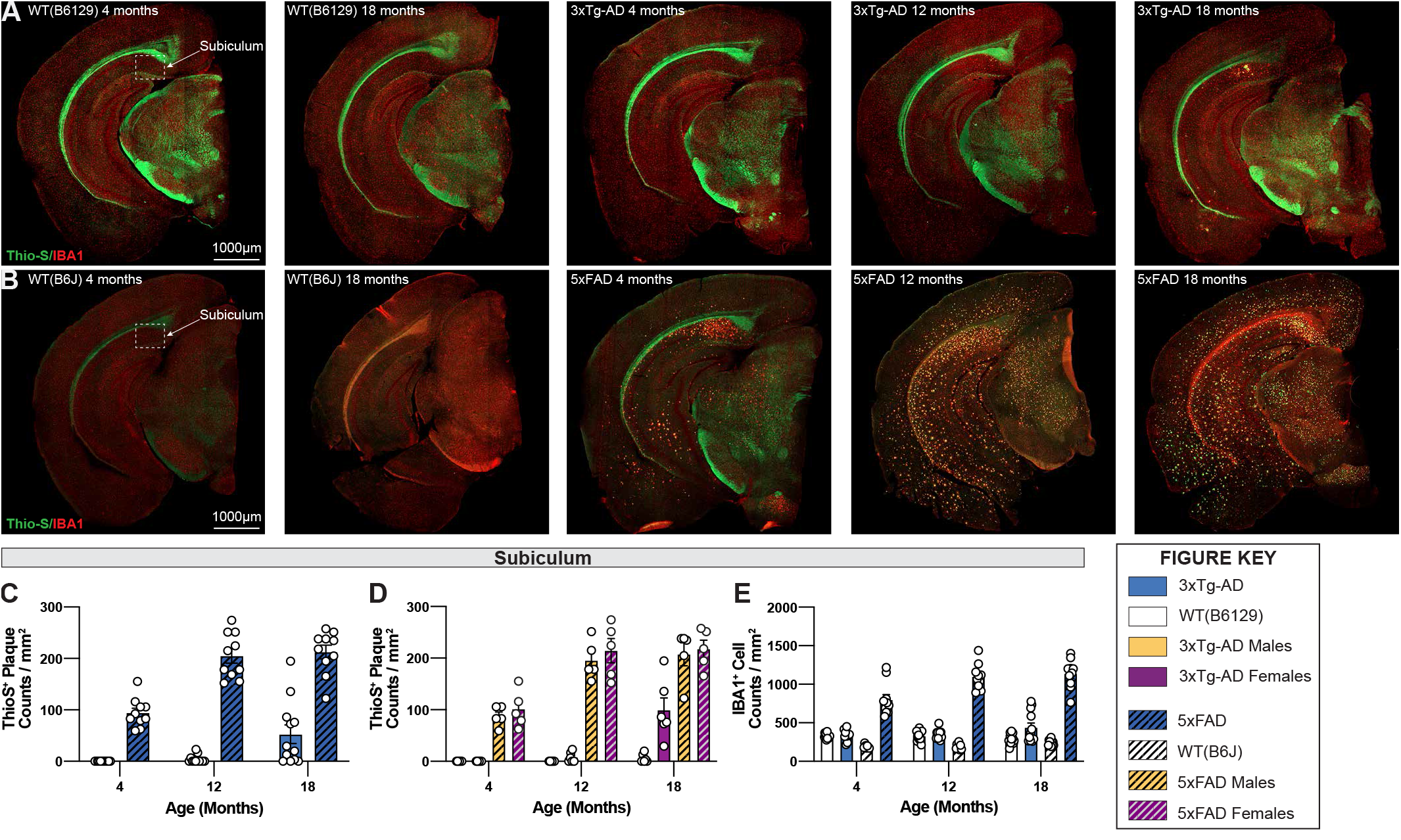
Comparison of fibrillar amyloid plaque accumulation in 3xTg-AD and 5xFAD mice. **(A)** Representative images of brain hemispheres of WT(B6129) at 4- and 18-month and 3xTg-AD mice at 4, 12, 18-month stained with Thio-S/IBA1. **(B)** Representative stitched brain hemispheres of WT(B6J) at 4- and 18-month and hemizygous 5xFAD mice at 4, 12, 18-month stained with Thio-S/IBA1. **(C-D)** Quantification for density of Thio-S^+^ plaques in subiculum hippocampal regions in 3xTg-AD and 5xFAD mice showed differences in plaque burden between mouse models and sexes. **(E)** Quantification for IBA1 immunostaining for microglia in subiculum hippocampal regions reveals age-related differences between mouse models. No stats n = 5-6 mice per genotype/age/sex. Data are represented as mean ± SEM.

## Discussion

We have evaluated the AD-related pathogenesis in current 3xTg-AD mice and highlight the drift that has occurred over the past two decades in terms of development of pathology. We show that 3xTg-AD mice develop both plaques and tangles in an age-related fashion, but that the timing of plaque development has been substantially retarded since the line was first produced. Also striking has been the emergence of profound sex differences in the development of both plaques and tangles, with these pathologies only emerging in female mice. Previously, we reported no sex differences in terms of pathology, but the development of deficits in female mice on stressful behavioral tasks (Clinton et al., 2007). Increased pathology in female mice is also seen in the 5xFAD mouse model, which uses the same Thy1 mini-gene regulating expression of the cDNA’s in the transgenes. The promoter in this Thy1 mini-gene was found to contain an estrogen response like element that can produce greater expression in females (Sadleir et al., 2015), which we recently replicated with our phenotyping efforts (Forner et al., 2021). Hence, the female sex-bias in pathology in both 3xTg-AD and 5xFAD AD models is likely due to increased expression from the Thy-1 mini-gene, rather than reflecting some inherent female-specific bias in AD susceptibility as found in the human population. Female 3xTg-AD mice develop both plaques and tangles, and their appearance coincides with loss of the extracellular matrix structures that surround mainly inhibitory neurons known as perineuronal nets, and the loss of parvalbumin+ interneurons. The fact that these pathologies develop slowly, and in an age-related fashion may be advantageous to studies that focus on late-onset AD and on the interactions between amyloid and tau pathologies. This finding contrasts with the aggressive amyloidosis models such as the 5xFAD mice, which produce magnitudes greater plaques from young ages (4+ months) and represent an excellent model for studying the effects of plaques on the brain (Oakley et al., 2006); Forner et al., 2021).

## Acknowledgements

This study was supported by the Model Organism Development and Evaluation for Late-onset Alzheimer’s Disease (MODEL-AD) consortium funded by the National Institute on Aging (U54 AG054349). The content is solely the responsibility of the authors and does not necessarily represent the official views of the National Institutes of Health.

